# Construction of relatedness matrices using genotyping-by-sequencing data

**DOI:** 10.1101/025379

**Authors:** Ken G Dodds, John C McEwan, Rudiger Brauning, Rayna M Anderson, Tracey C van Stijn, Theodor Kristjánsson, Shannon M Clarke

## Abstract

**Background:** Genotyping-by-sequencing (GBS) is becoming an attractive alternative to array-based methods for genotyping individuals for a large number of single nucleotide polymorphisms (SNPs). Costs can be lowered by reducing the mean sequencing depth, but this results in genotype calls of lower quality. A common analysis strategy is to filter SNPs to just those with sufficient depth, thereby greatly reducing the number of SNPs available. We investigate methods for estimating relatedness using GBS data, including results of low depth, using theoretical calculation, simulation and application to a real data set.

**Results:** We show that unbiased estimates of relatedness can be obtained by using only those SNPs with genotype calls in both individuals. The expected value of this estimator is independent of the SNP depth in each individual, under a model of genotype calling that includes the special case of the two alleles being read at random. In contrast, the estimator of self-relatedness does depend on the SNP depth, and we provide a modification to provide unbiased estimates of self-relatedness. We refer to these methods of estimation as kinship using GBS with depth adjustment (KGD). The estimators can be calculated using matrix methods, which allow efficient computation. Simulation results were consistent with the methods being unbiased, and suggest that the optimal sequencing depth is around 2-4 for relatedness between individuals and 5-10 for self-relatedness. Application to a real data set revealed that some SNP filtering may still be necessary, for the exclusion of SNPs which did not behave in a Mendelian fashion. A simple graphical method (a ‘fin plot’) is given to illustrate this issue and to guide filtering parameters.

**Conclusion:** We provide a method which gives unbiased estimates of relatedness, based on SNPs assayed by GBS, which accounts for the depth (including zero depth) of the genotype calls. This allows GBS to be applied at read depths which can be chosen to optimise the information obtained. SNPs with excess heterozygosity, often due to (partial) polyploidy or other duplications can be filtered based on a simple graphical method.

## Background

The calculation (via pedigree records) or estimation (using genotype data) of relatedness either implicitly or explicitly underlies most genetic analyses. Pedigree relatedness is used in the improvement of agricultural species and the elucidation of human diseases. More recently, marker-based estimates have been used to account for population structure in genome-wide association studies (GWAS) [1] and to enhance prediction of genetic merit [2] in agriculture through ‘genomic selection’ (GS). GS can be applied using an explicit estimate of the genomic relatedness matrix with genomic best linear unbiased prediction (GBLUP). Many other methods have also been proposed and although they might not explicitly use a relatedness matrix, they rely on phenotype similarity coinciding with marker similarity (i.e. relatedness) at specific locations on the genome.

The first applications of GWAS and GS have relied on genome-wide data generated using ‘SNP chips’ – assays that give genotype results for a pre-defined set of many thousands of single nucleotide polymorphisms (SNPs). An alternative method that is gaining popularity is to use genotype calls on the basis of next generation sequencing. This is usually implemented by sequencing a subset of the genome using restriction enzymes and size selection, so that more reads (i.e., higher read depth) of the sequenced regions can be obtained for a given sequencing effort [3]. The inference of genotypes from this method is often referred to as ‘genotyping-by-sequencing’ (GBS), although this term is also sometimes applied to deriving genotypes from whole genome sequencing. The application of GBS to plants has been reviewed by a number of authors [4-7] who discuss laboratory protocols, bioinformatics processes for genotype calling and the results of analyses using those genotypes. The various protocols allow different proportions of the genome to be sequenced [8] and therefore flexibility in the number of loci and the average read depth at an included genomic location. The subset of the genome that is sequenced is considered to be well spread throughout the genome; e.g., Poland *et al.* [7] conclude that GBS SNPs are approximately uniformly spaced on the physical genome (in wheat and barley). Therefore it is likely that much of the genome will be in linkage disequilibrium with SNPs assayed by this technique.

GBS is an attractive method for research in species for which SNP chips have not been developed, as it does not have the up-front development costs of the SNP chips. It may also be used in species without a reference genome sequence, although a reference genome is useful for sequence alignment, SNP ordering (if imputation or GWAS are to be used) and for quality control.

One drawback of GBS is that it is possible that only one of the two alleles is seen, and that a heterozygous individual is observed as homozygous. Also, if the coverage of the sequenced region is low, there will be many missing genotypes (much more than for SNP chips which commonly have call rates in excess of 99%). The usual analysis strategy is to filter the data so that most genotypes are called with sufficient depth to minimalize these errors and missing data (e.g. [6]). For a given cost, this results in fewer individuals being genotyped (to obtain greater depth per individual) or fewer SNPs being available (discarding those which do not pass the filtering step). However, Vela-Avitua *et al.* [9] find that genomic predictions based on identity by state methods benefit from having large numbers of SNPs, while Gorjanc *et al.* [10] find that genomic predictions using GBS benefit from including more individuals with many SNPs at relatively low depth. We investigate the implications of using GBS with low depth when estimating relatedness and suggest strategies for better relationship estimation. Our results apply to both full sequence and sequence from a random subset of the genome, but the latter is the more likely route due to its lower cost.

## Methods

### Methods of moments estimators for a single SNP

Suppose a SNP has alleles A and B. Let *g*\* denote the genotype as observed, e.g. AA* denotes that only A alleles are observed (for an individual), including the case where there is only a single read. Suppose

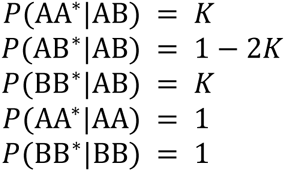

If allele reads are at random, and without error, then *K* = 1/2 ^*k*^ where k is the sequencing depth for the observation. For example, at *k* = ∞ *K* = 0 and the observed genotype is the true genotype. At *k* = 1, *K* = ½ and an AB genotype is seen as either AA or BB (each with probability ½). There may be other plausible models, for example ones where allele reads exhibit clustering such that observed homozygosity is higher (when depth exceeds one) than in the random reads case, but which still have *K* = ½ when *k* = 1.

Let *x* be the number of A alleles (the score) in the observed genotype (e.g. *x* = 2 for AA*). Subscripts *i* and *i’* will be used to denote (joint) genotypes or values relating to individuals *i* and *i’.* Let *p* be the A allele frequency (assumed known). Some commonly used estimates of relatedness are based on the quantities *S*_*ii*′_ = (*xi* − 2*p*) (*xi*′ − 2*p*) (for each SNP) [11]. We now derive expected values of these quantities (separately for the cases *i* = *i*’ and *i* ≠ *i*’).

If the individual has inbreeding *F*, then

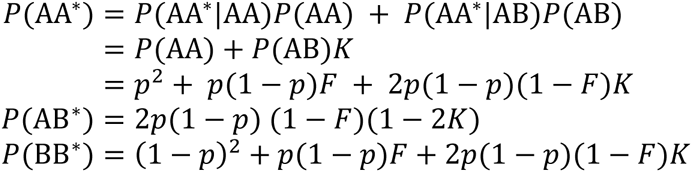

Joint observed genotypes can be written in terms of joint true genotypes (assuming that the values of *K*_*i*_ and *K*_*i*′_ are independent):

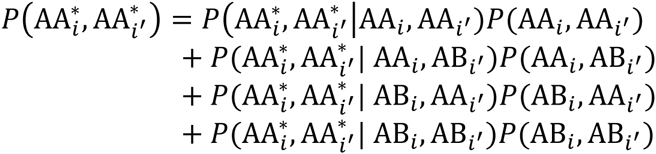

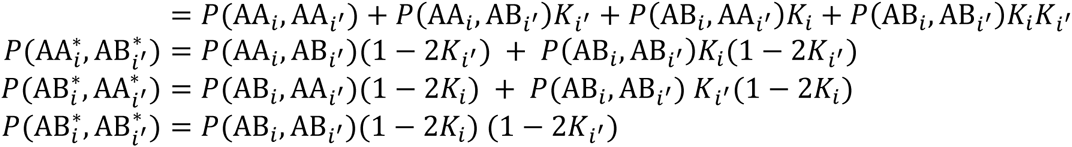

The probabilities of joint true genotypes can be written in terms of identity by descent measures (e.g. [12], p209). In our notation, the required probabilities for a biallelic locus are

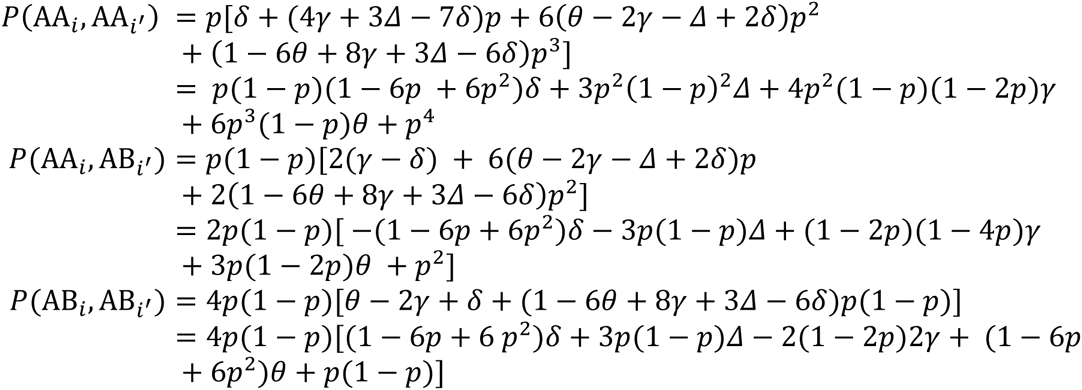

Further details about the quantities *θ*, *γ*, *δ* and *Δ* can be found in [12], but in particular, *θ* is the coancestry, also referred to as the kinship or half the relatedness, between two (different) individuals. We now derive expectations of relevant quantities.

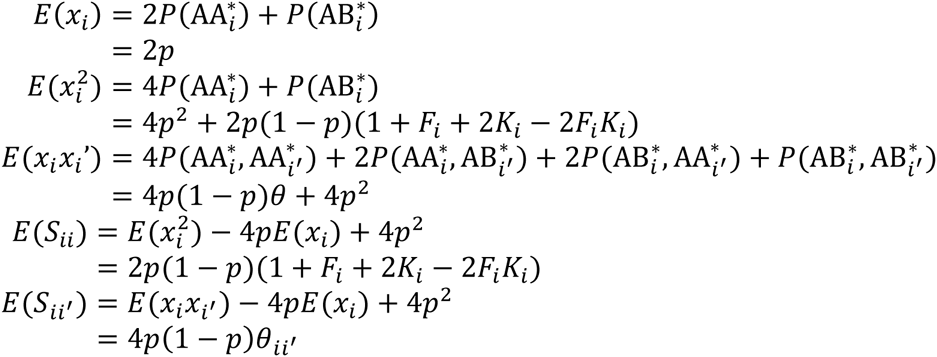

Notice that *E*(*S*_*ii*′_) does not depend on the *K*, i.e. on the sequencing depth. On the other hand, *E*(*S*_*ii*′_) is dependent on sequencing depth (unless *F* = 1 in which case the individual is completely homozygous); varying between 2*p*(1 − *p*)(1 + *F*) (for infinite depth) and 4*p*(1 − *p*) (for depth one). The latter result shows that *E*(*S*_*ii*_) increases with decreasing sampling depth – if sampling depth is not taken into account, the diagonals of the genomic relationship matrix will be inflated.

Rearranging the above equations, we obtain

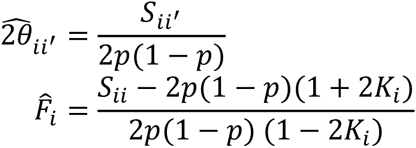

The divisor in the first of these equations is the one that is used (for both the diagonal and off-diagonals) when there are no depth issues for the genotype calls. It is seen to be the variance of the scores, and therefore the estimator is sometimes referred to as a correlation-based estimator. We prefer to regard it as a methods of moments estimator, with the denominator being chosen to make the estimator unbiased.

### Estimators using a sampled allele

We also consider an estimator which discards multiple reads for a SNP and individual, by randomly sampling one read from those available. Such an estimator may be more robust to model assumptions and/or be diagnostic for data that deviates from the model. However, when the model is correct, because the estimator does not use all the information, it will be more variable than one where all reads are used. The calculations are carried out as above, with *k* = 1, *K* = ½. For this estimator, *E*(*S*_*ii*_) = 4*p*(1 − *p*), i.e., it cannot be used to construct an estimator of *F*.

### Estimators using many SNPs

The above methodology relates to a single SNP. Let *j* index the SNP number, and where relevant (i.e., referring to an individual) the above quantities will be subscripted further by *i*. Let **Z** be the matrix (with number of rows equal to the number of individuals and number of columns equal to the number of SNPs) of centred genotype scores, i.e. having elements (*x*_*ij*_ − 2*p*_*j*_), where *i* indexes the rows (individuals) and *j* indexes the columns (SNPs). Then a genomic relationship matrix (GRM) can be calculated [11] as

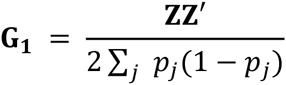

If the allele frequencies are considered as known, then from the above equations, off-diagonals of **G_1_** have expected value 2θ_ii’_, the relatedness between the individuals. Diagonal elements have expected values which depend on the *K*_*ij*_ (i.e. on the sampling depth at each SNP for that individual). To correct this, we replace the diagonal elements of **G** by

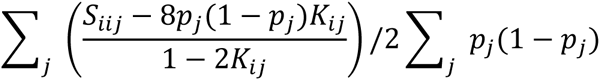

so that it is an estimator of 1 + *F*_*i*_ (the relatedness of an individual to itself). Note that if *K* = ½, the denominator in the first sum is zero. This relates to the case where *E*(*S*_*ii*_) does not depend on *F*, i.e., contains no information about *F*. Therefore, the two sums above are taken over the SNPs where *K* > ½.

A second estimator, **G**_3_, (we have reserved the use of ‘**G**_2_’ for 2nd GRM method of VanRaden [11], but do not use it in this article) uses the sampled alleles combined as for **G_1_**. As there is no information about self-relatedness, the diagonals of this GRM are ignored.

### Missing genotypes

If individuals are sequenced to a low depth, then many genotypes will be missing. Common approaches for dealing with missing genotypes are to replace them with estimated values, either using imputation [13] or more simply, by their population expected values (2*p*), sometimes referred to as ‘naïve imputation’ [14]. We propose an alternative strategy, based on the assumption that scored genotypes are a random sample across the genome. Elements of **G** are calculated using only those SNPs which are scored in both of the corresponding individuals. Here we denote the matrices constructed in this manner without and with the diagonal correction described above as **G_4_** and **G_5_** respectively. We refer to this fully corrected method of estimation (**G_5_**) as kinship using GBS with depth adjustment (KGD).

A drawback of this approach is that the set of SNPs in the calculation varies across relative pairs. An obvious calculation strategy would be to loop through each pair of individuals, take a copy of the genotype scores for those SNPs where both individuals are scored, and calculate the relatedness using those SNPs. However, software which enables matrix calculations is usually designed so that such computations are much faster than element-wise calculations. A strategy to take advantage of fast matrix computation (but that requires more memory) is as follows. Firstly, missing values in **Z** are replaced by zeros. This means that any SNP that is missing for either of a pair of individuals will not contribute to their corresponding element in **ZZ**′ (zero is added for that SNP). Next create a matrix **P0** with the same dimensions as **Z** with each row initially being identical and containing the allele frequencies, but then replace these by zero for individuals and SNPs with missing genotypes. Also create a matrix **P1** in the same way, but using one minus the allele frequencies. Then

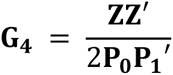

will only use SNPs where both individuals are scored, as required. The diagonals then need to be corrected, as above, to obtain **G_5_**. R software for undertaking these calculations is available from <u>https://github.com/AgResearch/KGD.git</u>

### Simulation

Several small simulations were undertaken to illustrate and compare methods. A set of 100 pairs of unrelated parents were generated along with two (full-sib) offspring of each pair of parents. A set of 10,000 SNPs were simulated with the allele frequencies sampled from a Uniform(0,1) distribution. The SNPs were modelled as being unlinked. Parent alleles were sampled from a population with these allele frequencies, while progeny received one of their parent’s alleles at random, and independently of the alleles received at the other SNPs (i.e., no linkage). Once the genotypes were created, GBS results were simulated by setting an average depth *d* (assumed constant over SNPs and individuals) and drawing *k*_*ij*_ alleles at random, with replacement, from the alleles of individual *i* at SNP *j*, where *k*_*i, j*_ ~ Poi(*d*) (i.e., a Poisson distribution with mean *d*). Simulations were undertaken for average depths 1, 2 and 8. Any SNP which had a minor allele frequency of zero in the genotype or GBS data was removed from the analysis. Relatedness estimates which required a value of *K* used the random alleles calculation, *K* = 1/2^*k*^ where *k* is the sequencing depth for the observation. Means and standard deviations (sds) of relatedness values were obtained over sets of full-sibs, parent-offspring, between each pair of individuals present as parents (unrelated) and selfs, for the various relationship matrix methods. We also calculate mean sample ‘call rates’ (the proportion of SNPs with at least one allele observed for that individual) and mean ‘co-call rates’ between pairs of individuals (the proportions of SNPs called in both individuals). Although the simulation of the full-sibs is not realistic in that it does not account for linkage, the aim here was to generate sets of individuals with a common level of relatedness, to allow a benchmark for comparing methods.

Another set of simulations were run to investigate the effect of sequencing depth on the sds of estimates, at a fixed total sequencing effort. The total effort was quantified as the number of SNPs times the mean depth. Total effort was fixed at 10,000 reads spanning a SNP and this effort was spread over between 500 and 40,000 SNPs (i.e. mean depth ranging from 0.25 to 20). For this simulation allele frequencies were sampled uniformly between 0.1 and 0.5 to ensure that all SNPs were retained in the analysis (i.e. to remove any simulation variability due to rare alleles being missing from the parents).

A third set of simulations investigated the effect of allele frequency distribution on the sds. For this simulation mean depth was fixed at 2 which gave near optimal results in the previous simulation. All SNPs were modelled as having the same allele frequency for a particular run. Each combination of allele frequency, ranging from 0.01 to 0.5, and number of SNPs, ranging from 1000 to 50,000, was simulated. Only the **G_5_** results are presented from these last two sets of simulations.

### Animals

The methods were also compared using a real dataset of 2203 Atlantic Salmon (*Salmo salar*) which were part of Stofnfiskur’s <u>(http://stofnfiskur.is/)</u> breeding programme. The dataset included fish from a cohort of 122 full-sib families, each having a unique dam, but with 66 different sires each siring 1-3 families. Genotypes were obtained from all sires, 119 dams and 94 full-sib families (range 13-37 and average 18.9 progeny per family for a total of 1773 progeny). In addition, there were a set of 244 ‘unrelated’ fish (sequenced one to four times, average 2.5), plus an ‘unrelated’ reference which was sequenced 27 times. Each time a sample was sequenced it consisted of approximately 2 million 100bp reads (see below) and these sampled approximately 1.2% of the genome or 30Mbp to an average depth of 5 fold.

### Genotyping

Tissue samples for DNA extraction were collected in the form of a fin clip from each animal which was stored in 96% ethanol. A subsample of approximately 3 mm 2 was taken from each fin clip and stored in a 96 deep well plate along with 50 µl of 96% ethanol until required. Prior to extraction, the tissue was air dried overnight to remove all traces of ethanol and then the DNA was extracted following the high throughput tissue extraction method as described by Clarke *et al.* [15]. The amount and quality of the DNA was first examined via a Nanodrop 8000 (Thermo Scientific, Weltham, Massachusetts, United States) spectrophotometer. A subset of the extracted DNA samples were also visually assessed via 1.0% agarose gel to ensure high molecular weight DNA was present. Picogreen (Quant-iTTM Picogreen® dsDNA Reagent, Cat P11495, Life Technologies, Carlsbad, California, United States) fluorescence was utilised to accurately quantify the DNA prior to generating the restriction enzyme digest fragment sequencing libraries. These GBS-libraries were prepared utilising the *Pst*I restriction enzyme following the method outlined in Elshire *et al.* [3]. The oligonucleotides to form the barcode adapters and common adapters with *Pst*I overhangs were purchased from MWG Operon (www.operon.com) and the 96 barcodes were designed by Deena Bioinformatics (<u>http://www.deenabio.com/services/gbs-adapters</u>). The adapters were prepared and combined with 100 ng of DNA sample as outlined in Elshire *et al.* [3]. Subsequent GBS-library preparation was also carried out as described in Elshire *et al.* [3] with the following exceptions. The *Pst*I digest was performed as follows: samples (DNA together with adapters) were digested with 5 U *Pst*I (New England Biolabs, Ipswich, Massachusetts, United States) in 20 µl volume containing 1x NEB CutSmart buffer and incubated at 37 ^°^ C for 2 h followed by heat inactivation (65 ° C for 30 min). Following digestion, ligation was carried out by adding a 30 µl ligase solution (containing 5 µl ligase buffer with ATP and 1 µl T4 ligase (NEB) and incubating at 22° for 1 h followed by heat inactivation (65 ° C for 30 min). The individual barcoded samples were combined (5 µl of each), purified using a commercial kit (QIAquick PCR Purification Kit; Qiagen, Valencia, CA) according to the manufacturer’s instructions and eluted to a final volume of 50 µl. The restriction fragment library was then amplified in 4 individual reactions (total volume of 50 µl, with 4 µl pooled barcoded DNA fragments, 1x *Taq* Master Mix (NEB) and 25 pmol of each primer), at 72 ° for 5 min, 98 ° for 30 s followed by 18 cycles of 98 ° for 30 s, 65 ° for 10 s, 72 ° for 30 s and a final Taq extension step at 72 ° for 5 min. The amplified sample pools were then combined, purified as above, eluting in 35µl of EB Buffer, and then further purified utilising a Pippin Prep (SAGE Science, Beverly, Massachusetts, United States) to select the DNA sequencing library in the size range of 150-500bp. The library was eluted in 40µl of the supplied buffer and a 1 µl aliquot of a 1:5 dilution was evaluated on a 2100 Bioanalyser (Agilent Technologies, Santa Clara, California, United States) to establish the quality of the library. The concentration was determined by quantifying 1µl of library on a Qubit Fluorometer (Life Technologies, Carlsbad, California, United States). Single-end sequencing (1x100) was performed on an Illumina HiSeq2500 utilising v4 chemistry, yielding approximately 25Gb of raw sequence data per lane.

Raw fastq files were quality checked using FastQC v0.10.1 (<u>http://www.bioinformatics.babraham.ac.uk/projects/fastqc/</u>). For each lane a random subsample of 10,000 reads was checked for contamination. The subsamples went through adapter removal using cutadapt [16] with setting -a AGATCGGAAGAGCGGTTCAGCAGGAATGCCGAGACCGATCTCGTATGCCGTCTTCTGCTT, then mapping onto the *Salmo salar* reference genome (Ssa_ASM_3.6, <u>http://www.icisb.org/sequence</u>) with bwa version 0.7.9a-r786 [17] with default settings, except bwa aln -B 10.

Between 124k and 51M raw reads were processed with UNEAK, Tassel version 3.0.170, [18] to detect variants and report reference and alternative allele counts at variant sites. UNEAK settings used were (1) -UFastqToTagCountPlugin -c 1 -e PstI,

(2) -UMergeTaxaTagCountPlugin -m 600000000 -x 100000000 -c 12,

(3) -UTagCountToTagPairPlugin -e 0.03, (4) -UMapInfoToHapMapPlugin –mnMAF 0.03 -mxMAF 0.5 -mnC 0.1.

### Analysis

The allele count data was analysed using the relatedness estimation methods discussed above. Allele frequencies were estimated using the total allele counts from all reads. A number of additional quality control (QC) diagnostics were also undertaken, and the data filtered on the basis of these results, with relatedness estimation applied both pre-filtering and post-filtering using a variety of filters. Samples were examined for sequence depth. The SNP ‘call rates’ (the proportions of individuals with at least one sequence read at each SNP position) and their minor allele frequencies (MAFs), based on genotype calls, were calculated. Hardy-Weinberg disequilibrium (observed frequency of the reference allele homozygote minus its expected value) was plotted against MAF, using a colour gradient to illustrate SNP average depth. We refer to this as a ‘fin plot’ because of the shape of the boundaries (upper bound of MAF-MAF^2^, lower bound of –MAF^2^). The recorded pedigree was used to assess the effectiveness of the QC steps and of the various relatedness estimates. Finally, parentages which were not confirmed by the GBS results (using the final chosen filtering steps) were assumed unknown.

## Results

### Simulation

Mean relatedness estimates for the different relationship sets are shown in Figure 1. Many of the method and scenario combinations produced very similar means near the true values of 1 (for selfs), 0.5 (for both full-sibs and parent-offspring) and 0 (unrelated parents). This includes all the results (labelled ‘Chip’) using the true simulated genotypes (with **G_1_**) and the methods proposed here for use with GBS data (**G**_3_ and **G_5_**). All methods gave estimates close to zero for the unrelated.

**Figure 1.**
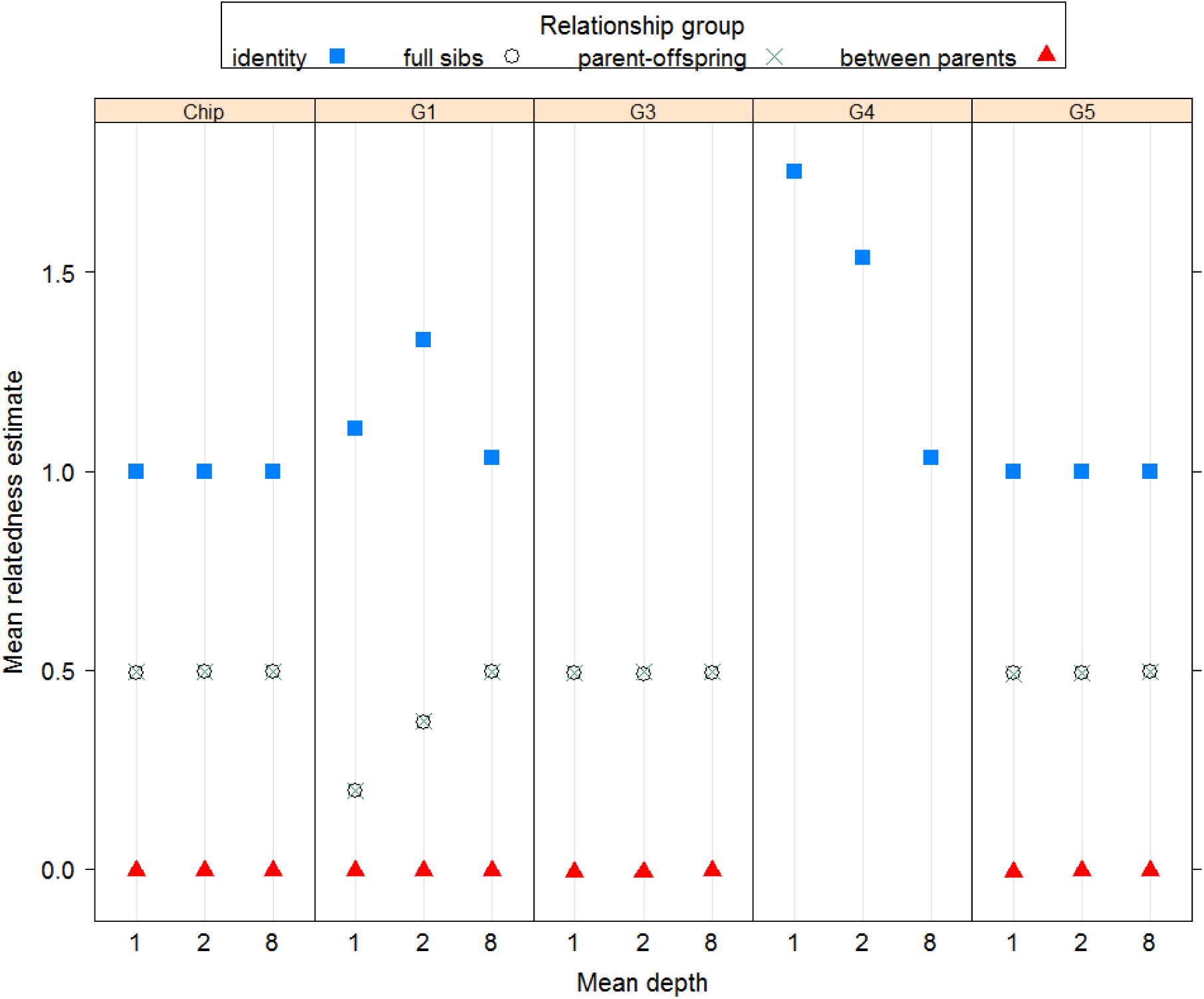
Mean relatedness estimates in simulated data using different methods. Mean estimates of relatedness for each individual with itself (identity), all pairs of full-sibs, all parent-offspring pairs, and between parents. Each estimation method is shown in a separate panel. Estimates using **G_1_** with the actual genotype data are shown as ‘Chip’. The other panels show estimates using the different **G** matrix methods with simulated GBS data. **G_4_** results are only shown for the identity group (the method is the same as **G_5_** for relatedness between individuals). The different sets of ‘Chip’ results correspond to the different sets of GBS simulations (at different depths).

Use of **G_1_** with GBS data gave downwardly biased estimates for pairs of related individuals (due to the inflated denominator). This bias decreased with sequencing depth and had essentially disappeared when *d* = 8. Use of **G_1_** with GBS data gave upwardly biased estimates for self-relatedness. The bias is due to an inflated numerator (not accounting for depth) which appears to exceed the effect of the inflated denominator. The most bias was seen at *d* = 2 and the least at *d* = 8. The use of method **G_4_** removes the inflation in the denominator, and so results in more upwardly biased estimates for self-relatedness than did the use of **G_1_**. Additionally correcting for sample depth in the estimates for self-relatedness (method **G_5_**) gives results very close to true values for all the relationship types.

The standard deviations (sds) of the relatedness estimates for the different relationship sets are shown in Figure 2. The sds given here are across pairs of individuals, so will include any variation in relatedness in that set of pairs. For the relationships examined, only full-sibs have variation in their true relatedness, but this will be very minor here, because the SNPs have been simulated independently. Using the true simulated genotypes (with **G_1_**; labelled ‘Chip’) gave the lowest sds, while all the GBS methods except **G**3 gave similarly low sds when *d* = 8. Among the GBS methods, **G**3 gave the highest sds, followed by **G_5_**. However, these were the only two methods which gave unbiased mean estimates. Their higher sds reflect that they correctly account for the information in the GBS data. For these two methods the sds decreased with increasing mean depth. However the sds for **G**3 with higher depth did not approach those for genotype data (shown as ‘Chip’ in Figure 2) because increasing depth only increases call rate and co-call rate for an individual or a pair, i.e. the method never uses the full information at a SNP, in contrast to **G_5_** which is effectively using the genotypes when depth is high.

**Figure 2.**
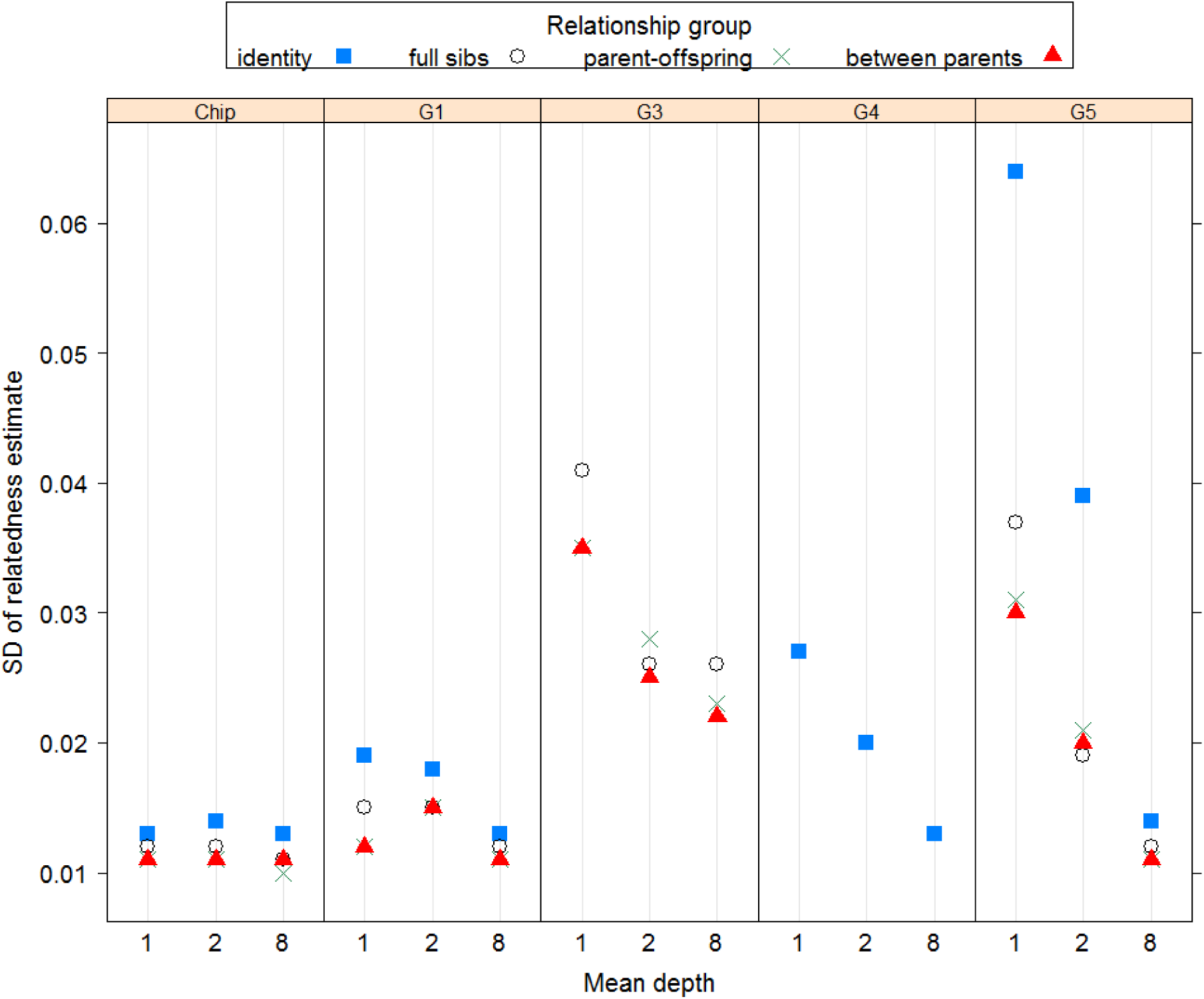
Standard deviations of relatedness estimates in simulated data using different methods. Standard deviations of estimates of relatedness for each individual with itself (identity), all pairs of full-sibs, all parent-offspring pairs, and between parents. Each estimation method is shown in a separate panel. Results using **G_1_** with the actual genotype data are shown as ‘Chip’. The other panels show results using the different **G** matrix methods with simulated GBS data. **G_4_** results are only shown for the identity group. The different sets of ‘Chip’ results correspond to the different sets of GBS simulations (at different depths).

Figure 3 shows the relationship between means and sds of estimated relatedness and the mean depth of each SNP when the total sequencing effort (number of SNPs times mean depth) is fixed at 10,000. The mean relatedness values show a small deviation from their expected values at very low mean depth. This is possibly a consequence of having estimated allele frequencies from the sequence reads. At low depth, there will be fewer individuals scored for any particular SNP. For example, at mean depth 0.25 each SNP was scored in 44 (of 200) parents, on average. By the time mean depth reached 1, each SNP was scored in 127 parents on average.

**Figure 3.**
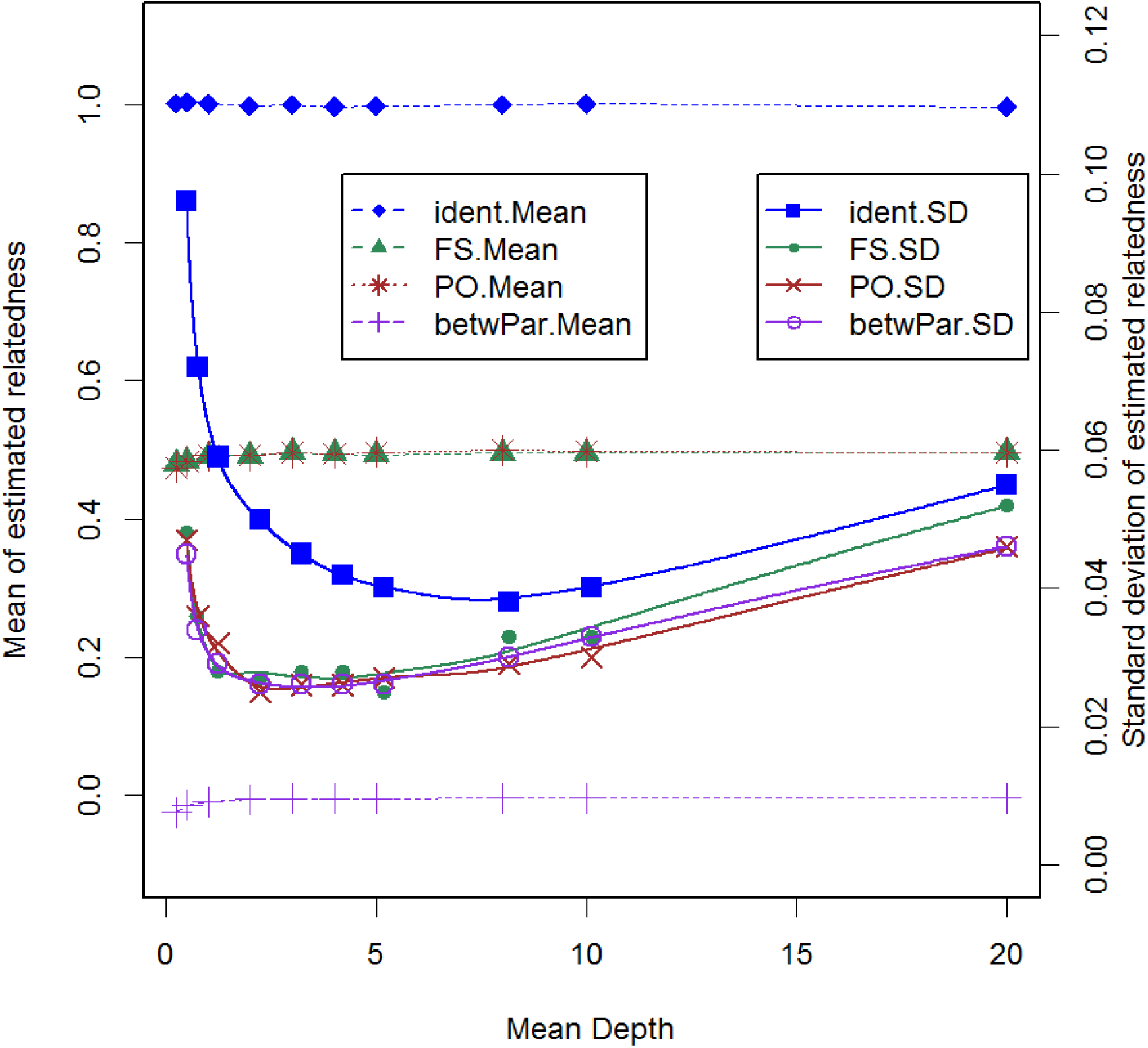
Means and standard deviations of relatedness estimates with constant sequencing effort. Means (.Mean) and standard deviations (.SD) of **G_5_** estimates of relatedness for each individual with itself (ident), all pairs of full-sibs (FS), all parent-offspring pairs (PO), and between parents (betwPar) at different mean depths, with the total sequencing effort held constant at 10,000 reads.

The sds of estimated relatedness initially decline rapidly with increasing mean depth, and then slowly increase. For relatedness for pairs of individuals, the minimum (smoothed) values are around a depth of 2 to 4. For self-relatedness the minimum value is near a depth of 8, although any depth between 4 and 10 gave similar results. For self-relatedness, greater depth is needed to ensure a reasonable proportion of SNPs have depth of at least 2, which is the minimum required to allow an estimate of inbreeding.

A contour plot of standard deviations of estimates of relatedness for parent-offspring pairs (true relatedness exactly 0.5) at a mean SNP depth of 2 is shown in **Figure 4**. It shows that around 30,000 SNPs with allele frequency 0.01 would be needed to obtain the same precision as around 1100 SNPs with allele frequency 0.5. Similar results were found for full-sibs (not shown). Standard deviations of relatedness estimates between parents appeared almost independent of the allele frequency, while for self-relatedness 20,000 SNPs with allele frequency 0.05 had similar precision to 1000 SNPs with allele frequency 0.3 (not shown).

**Figure 4.**
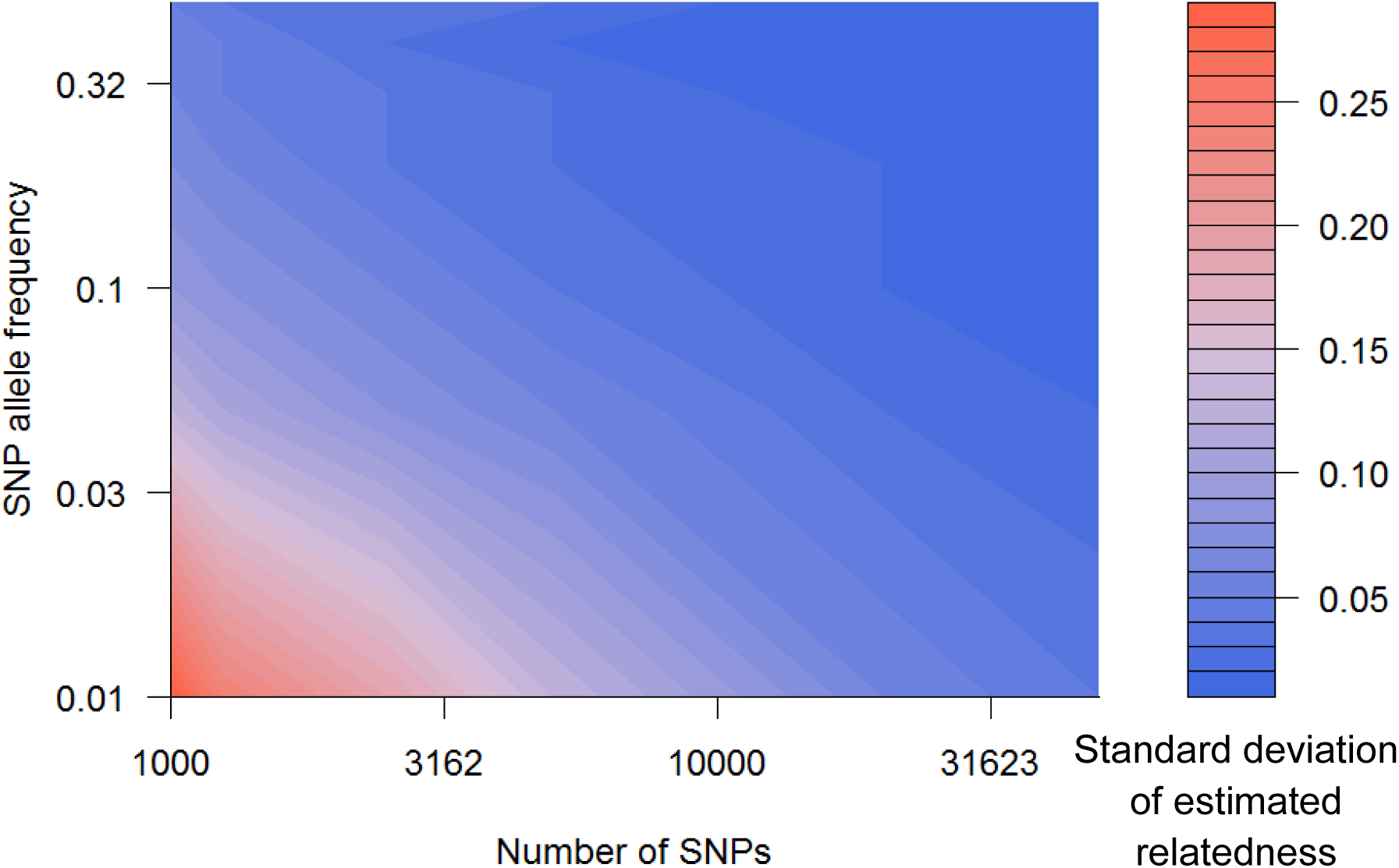
Standard deviations of relatedness estimates by numbers of SNPs and their allele frequencies. Contour plot of standard deviations of **G_5_** estimates of relatedness for all parent-offspring pairs when using sets of SNPs with the same allele frequency, for different allele frequencies and different numbers of SNPs, sequenced to a mean depth of 2. Axes are on the log scale.

The mean depth (2) used in this set of simulations is not optimal for self-relatedness (standard deviations about 30% higher than at the optimal depth), but a similar relationship between allele frequency and number of SNPs would be expected at higher depths.

### Atlantic Salmon results

The sequencing and conservative bioinformatics process, typical of what would be expected using a species with no available genome sequence, resulted in 30,923 SNPs for analysis. The mean SNP depth was 7.9 while 34% of animal by SNP combinations had no results. The mean depths ranged from 0.38 to 38.10 for each animal, except for the reference (mean depth of 89.06). Although there was a wide range (partly due to some animals being run more than once), none appeared to be ‘outliers’, so all were retained for further analysis. The distribution of minor allele frequencies (MAFs) is shown in Figure 5 and shows that the highest density of SNPs had high MAF (near 0.5), while low MAF (close to the 0.03 cut-off set in the bioinformatics process) SNPs were also common.

**Figure 5.**
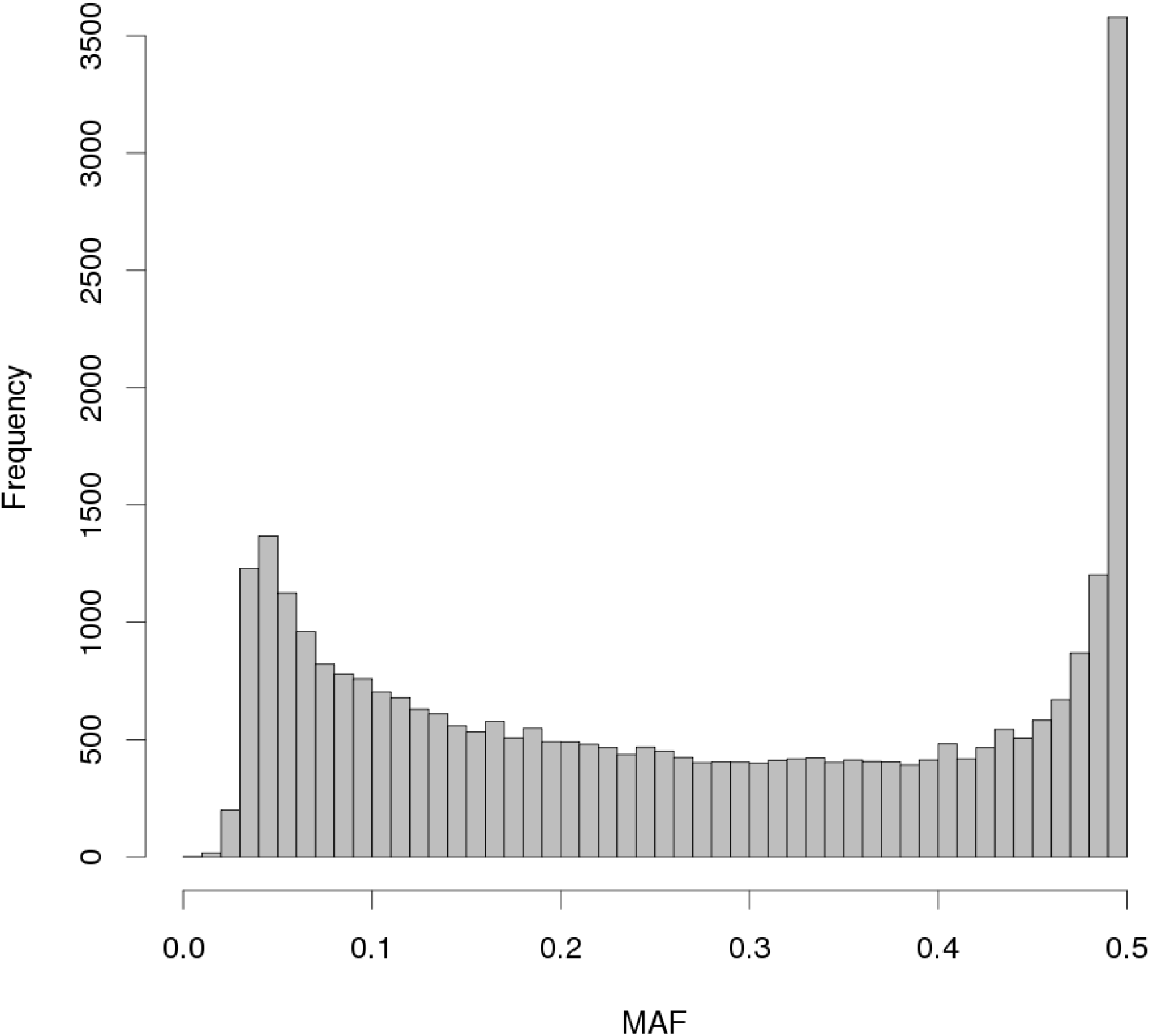
MAFs of the salmon data. Distribution of minor allele frequencies (MAF) for the Atlantic salmon data.

**G_5_** estimates of relatedness are shown in Table 1. The initial analysis (‘All’) did not filter the data after the bioinformatics step. Estimates for pairs of related animals were lower than their pedigree relatedness (i.e. lower than 1 for identity, and lower than 0.5 for both full-sibs and for parent-offspring). Estimates for pairs from the offspring cohort having different parents were higher than their pedigree relatedness of 0. This latter result could be due in part to relatedness among parents as the pedigree provided did not show ancestors further back than parents. Here we use pedigree relatedness as the gold standard, but note that there may be errors in pedigree recording, which would tend to make the full-sib and parent-offspring sets less related, on average, than that given by their recorded pedigree.

**Table 1.**
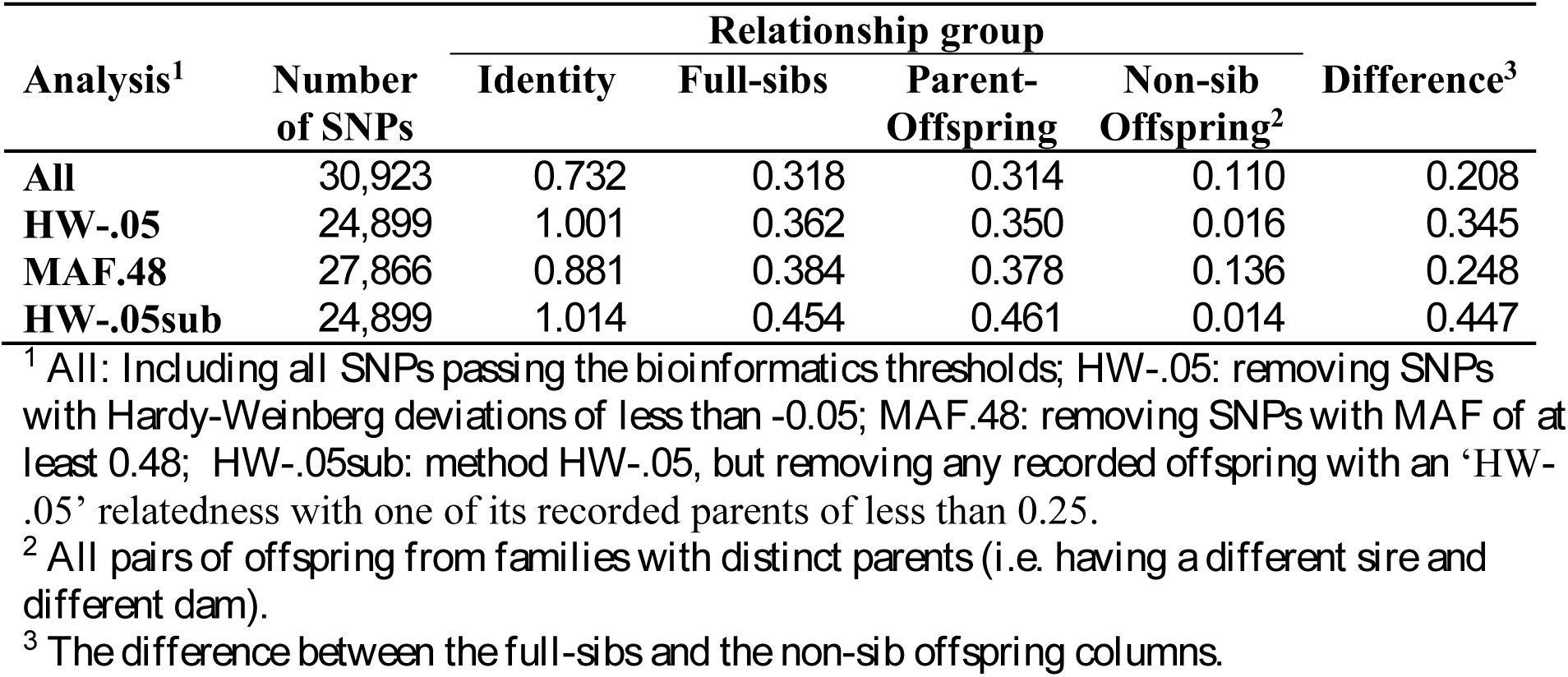
Mean relatedness estimates. Mean **G_5_** relatedness estimates for animals with the same pedigree relatedness. <tbl/>

| Analysis<sup>1</sup> | Number of SNPs | Relationship group |  |  |  | Difference<sup>3</sup> |
| --- | --- | --- | --- | --- | --- | --- |
|  |  | Identity | Full-sibs | Parent-Offspring | Non-sib Offspring<sup>2</sup> |  |
| **All** | 30,923 | 0.732 | 0.318 | 0.314 | 0.110 | 0.208 |
| **HW-.05** | 24,899 | 1.001 | 0.362 | 0.350 | 0.016 | 0.345 |
| **MAF.48** | 27,866 | 0.881 | 0.384 | 0.378 | 0.136 | 0.248 |
| **HW-.05sub** | 24,899 | 1.014 | 0.454 | 0.461 | 0.014 | 0.447 |

<sup>1</sup> All: Including all SNPs passing the bioinformatics thresholds; HW-.05: removing SNPs with Hardy-Weinberg deviations of less than -0.05; MAF.48: removing SNPs with MAF of at least 0.48; HW-.05sub: method HW-.05, but removing any recorded offspring with an 'HW-.05' relatedness with one of its recorded parents of less than 0.25.
<sup>2</sup> All pairs of offspring from families with distinct parents (i.e. having a different sire and different dam).
<sup>3</sup> The difference between the full-sibs and the non-sib offspring columns.

The diagonals of **G_5_** (self-relatedness estimates) were plotted against the log-transformed mean depth for each sample as a diagnostic for the effectiveness of the depth adjustment used in **G_5_**. This is shown for the ‘All’ analysis in Figure 6, and shows that estimates of self-relatedness increase with sequencing depth, an unexpected feature.

**Figure 6.**
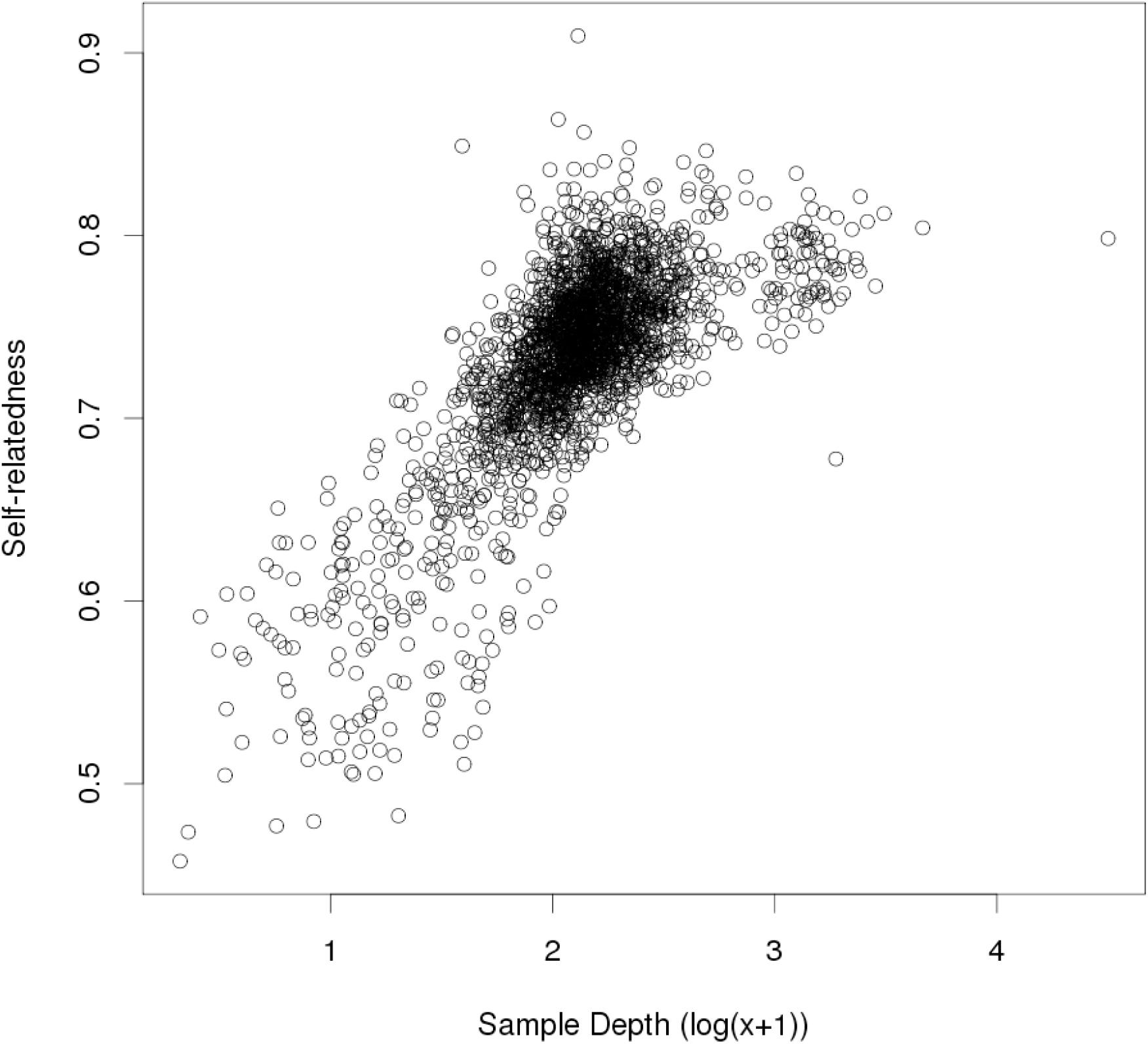
Self-relatedness estimates without SNP filtering. **G_5_** estimates of self-relatedness, from the analysis with no SNP filtering (‘All’), plotted against log-transformed sample depth.

In view of these unexpected results, further diagnostics were undertaken, including a ‘fin plot’ (Figure 7). The upper edge of this plot has a high density of low depth (coloured grey) SNPs. These will have a depth of 1 in many individuals, in which case the corresponding observed genotype is a homozygote, and therefore they appear near the upper boundary of maximum homozygosity (at the given MAF). There is a region of intermediate depth SNPs near the middle of the plot (i.e. disequilibrium near zero). Finally there are many high depth (coloured blue) near the lower boundary, i.e., with maximal heterozygosity (at the given MAF), and many of these have high MAF. It is quite likely that the latter group represents calls from regions of genome duplications or repetitive regions. These regions would tend to have at least double the apparent depth than SNPs from the other regions. Atlantic salmon has been found to have many such regions [19]. Any putative SNPs in these regions will not follow Mendelian (diploid) inheritance, as assumed in the development of the relatedness estimates used here.

**Figure 7.**
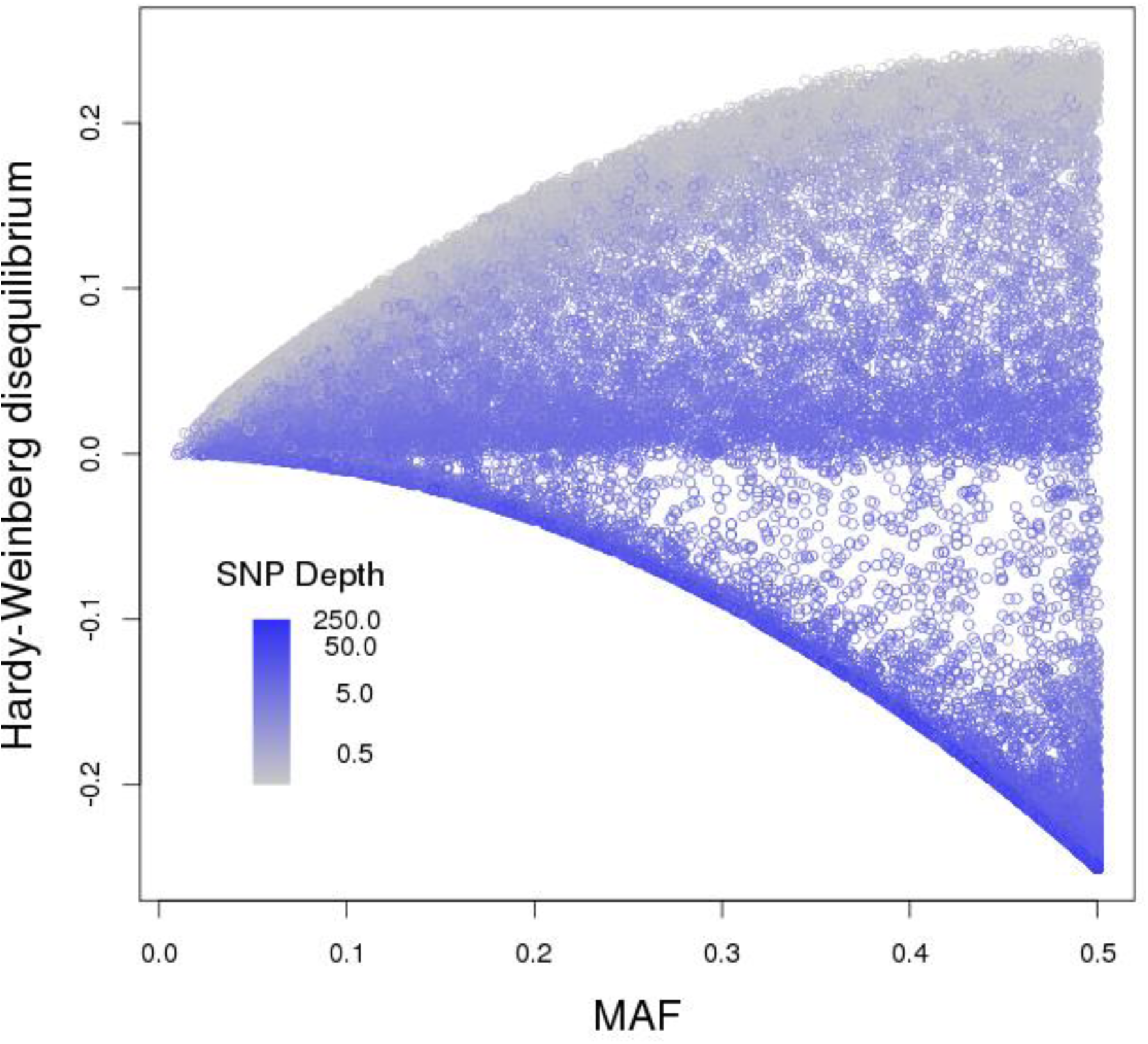
Fin plot for all SNPs. ‘Fin plot’ for analysis ‘All’. Hardy-Weinberg disequilibrium is plotted against MAF. Points are shaded from grey (low SNP depth) to blue (high SNP depth).

We have tried various filters based on this plot, such as proportional closeness to the lower boundary or a cut-off based on level of Hardy-Weinberg disequilibrium. The best results (in terms of giving relatedness estimates the closest to the pedigree-based values) of those investigated was to remove SNPs with Hardy-Weinberg disequilibrium below -0.05 (although a cut-off of -0.10 was only marginally worse). These results are shown in Table 1 as analysis ‘HW-.05’. This filter has improved the estimated relatedness for all groups, with higher values than the ‘All’ analysis for identity, full-sibs and parent-offspring pairs, and lower estimated relatedness for non-sib offspring pairs. The filter has also removed the relationship between the estimate of self-relatedness and the mean depth for the animal (Figure 8). The mean depth for the filtered SNPs was 3.3.

**Figure 8.**
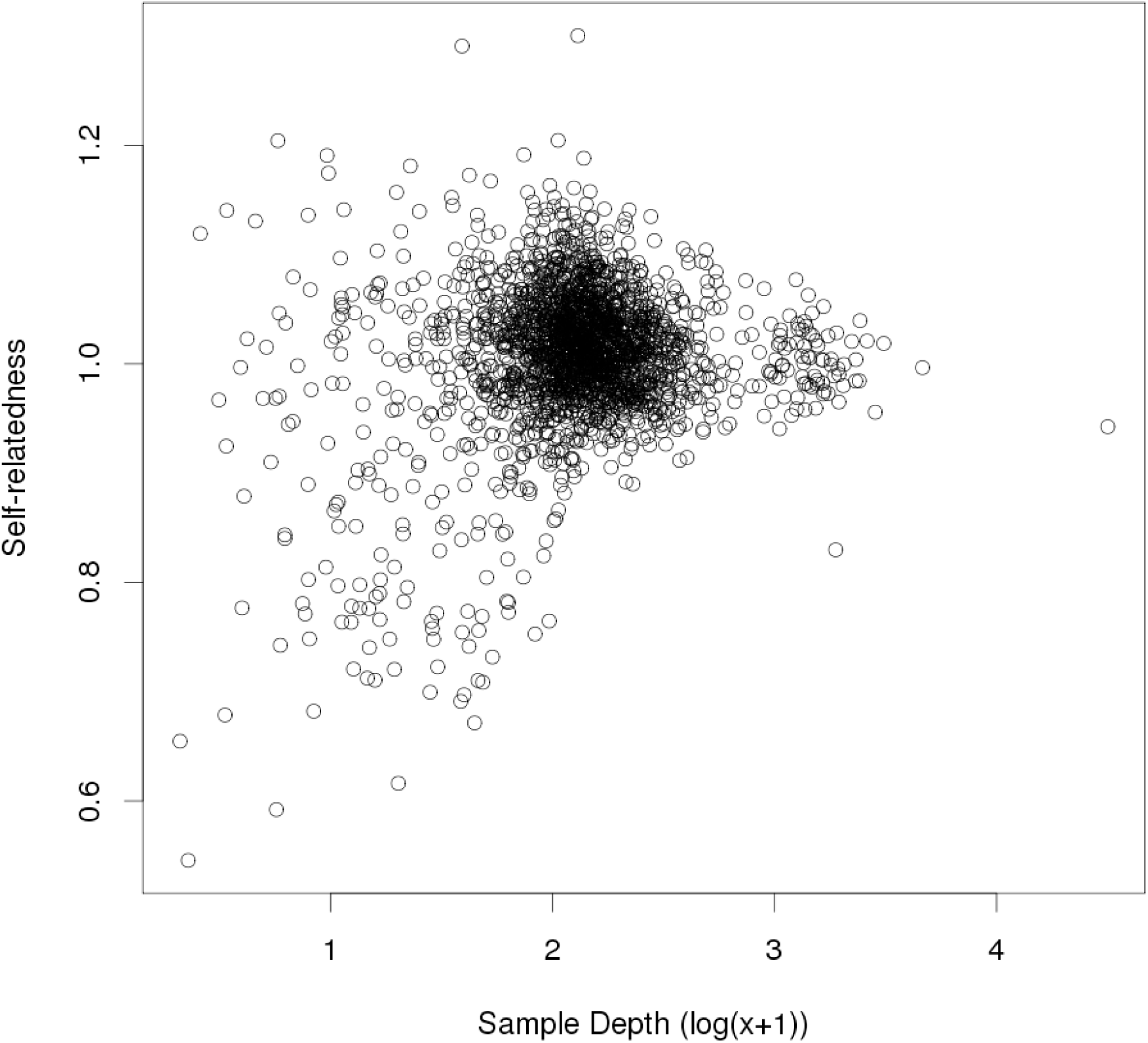
Self-relatedness estimates after SNP filtering. **G_5_** estimates of self-relatedness, from the analysis with SNPs having Hardy-Weinberg disequilibrium less than -0.05 removed (Analysis ‘HW-0.05’), plotted against log-transformed sample depth.

Although the plot of MAF suggests that it is only high MAF SNPs that may be causing a problem, a filter discarding SNPs with MAF≥0.48 (Analysis ‘MAF.48’) did not perform as well as HW-.05 (Table 1). Although this filter improved estimates for full-sibs and parent-offspring pairs, it had higher estimates for non-sib offspring, and identity estimates were lower than for HW-.05 and still showed a relationship with SNP depth (data not shown).

Even after filtering, the relatedness estimated between related pairs was lower than expected. Examination of the distribution of relatedness estimates for parent-offspring pairs, using ‘HW-.05’ showed a bimodal distribution, with peaks near 0 and 0.5. This suggests that there have been some errors in assignment (possibly parentage recording and/or sample tracking). The relatedness statistics were recalculated after removing any recorded offspring with an ‘HW-.05’ relatedness with one of its recorded parents of less than 0.25. These are shown in Table 1 as ‘HW-.05sub’. This resulted in relatedness values close to the expected 0.5 (or higher) for parent-offspring and for full sib pairs, compared with the full set of animals (HW-.05); estimates for other relationships remained at similar values.

### Discussion

The cost of sequencing has declined rapidly, especially with the introduction of ‘next generation’ sequencing technology. For many applications, full genome sequencing of an individual, to obtain SNP genotypes, is still not cost-effective. A promising strategy is to use ‘genotyping-by-sequencing’ (GBS) which allows a subset of the genome to be sequenced [3]. The technology has been designed for sequencing a subset of the genome of individuals, but DNA barcoding allows samples to be combined, and therefore yielding, for the same sequencing effort, similar sequencing depth to that obtained from a full genome run. GBS is particularly attractive for species for which other high-throughput technologies (such as SNP arrays) have not been developed.

GBS has already been investigated for genomic prediction in a number of species, such as wheat [20], *Drosophila,* using whole genome sequencing, [21], maize [22], soybean [14], spruce [23] and ryegrass [24]. These studies use a range of mean depth (from approximately 1, [22], to greater than 10, e.g. [14]). Often the data is filtered (e.g. [6]), where SNPs below a particular call rate are discarded, while missing genotypes are imputed using a range of algorithms, including naïve imputation. Some studies compared imputation methods, but generally found little difference in terms of genetic value prediction accuracy (e.g. [20]). Crossa *et al.* [22] included a ‘nonimputed’ comparison (naïve imputation with the weighting adjustment for missingness given by VanRaden [11]) and found it similar to results with a more sophisticated imputation method. Most studies used GBLUP as the prediction method, but some studies also included one or more other prediction methods. In general there was little difference between methods in prediction accuracy.

It is unclear whether these studies would find improved accuracies using the methods recommended here. Given that alternative prediction methods gave similar accuracies, it is possible that these other methods suffer from similar issues as when using a relationship matrix (via GBLUP), but that the effects of (e.g.) naïve imputation and not adjusting self-relatedness for depth are indirect. Alternatively, the corrections we propose here may not affect prediction accuracy unduly. This is supported by the simulation study of Gorjanc *et al.* [10], who found that (using a ridge regression model, which can be equivalent to GBLUP [25]) with a large number of SNPs (at least 60,000) genotyped by GBS, genomic prediction accuracy did not decline until the mean SNP depth dropped below 1. We have shown that mean SNP depths of 1-2 have important effects on the GRM **G_1_** (calculated without adjustments), but this does not appear to have affected prediction accuracy in [10]. Nevertheless it would seem sensible to make the adjustments proposed here, so that the GRM correctly reflects the relatedness. Further research is needed to understand whether similar adjustments can be made for other prediction methods.

Although naïve imputation leads to downwardly biased estimates of relatedness, it is not clear whether more sophisticated imputation methods will suffer from the same issue. It would seem likely that the better the imputation is at recovering the true genotype, the less the bias in relatedness estimates, but this needs further investigation. An attraction of GBS is that it can be applied with minimal genomic information, so imputation methods that require knowledge of SNP order may not be an option. Nevertheless, imputation methods that improve on naïve imputation are being developed [26, 27].

When relatedness estimation is used only for data auditing, the focus is on a pair of individuals, and calculations are based on only those SNPs scored in both individuals [28]. We have taken this approach to dealing with missing genotypes, and have shown that naïve methods can produce severely biased estimates. VanRaden [11] proposed an adjustment to the GRM to account for missing SNPs, and this was applied in the study of Crossa *et al.* [22], as discussed above. This applies a weight to each individual based on which SNPs are missing, so is not specific to the pair of individuals under consideration. We applied this method in our first simulation (comparing GRM methods) for the case with mean depth 1. We obtained exactly the same result for self-relatedness as method **G_4_** (which adjusts for missing SNPs, but not SNP depth). For self-relatedness there is only missingness for one individual to consider. However, the method only partly improved estimates of relatedness for full-sibs and parent-offspring groups (mean relatedness estimate of 0.31 for both groups). Other researchers have also noticed issues with GRM constructed from GBS data. Pérez-Enciso [29] found that estimates were ‘distorted’ in GRM from low depth (4) simulations.Ashraf *et al.* [30] and Cericola *et al.* [24] noted inflated diagonals and shrunken off-diagonals and have proposed adjustments. Ashraf *et al.* [30] also noted that the inflated diagonals led to inflated estimates of genetic variance. Gorjanc *et al.* [10] found inflated or shrunken estimates of genetic merit (depending on the number of SNPs in training and validation) and that these effects were more pronounced at low coverage. This was despite fixing genetic and residual variances at their true values.

We have considered the estimation of relatedness using a popular method (the **G_1_** GRM). Theoretical investigations showed that an adjustment was necessary for the diagonals of the GRM. We also argued that the divisor of each element in the GRM should only use SNPs that are in the corresponding calculation for the numerator. The adjustments were corroborated by a small simulation, where estimates of relatedness appeared to be unbiased, except at very low depth. The downward bias at low depth (Figure 3) could be due to poorer estimation of allele frequencies (due to a lower number of observations for each SNP) – this was supported by a further simulation using the true value of the allele frequency in the calculations, which showed about a 10 or 50-fold reduction in bias for parent-offspring and full-sibs, respectively. Our calculations have assumed that allele frequencies were known, while in practice they must be estimated, often from the same data. Allele frequency estimates based on larger samples are preferred, and for this reason we included the available non-family fish as well as the family sets when estimating allele frequencies. These allele frequencies were based on allele counts, rather than genotype counts (equal weighting for genotyped individuals). In this analysis these options gave similar results, but appropriate weighting for genotypes based on variable depths is a topic for further research. The issue of appropriate allele frequencies has been discussed by Powell *et al.* [31] and Makgahlela *et al.* [32].

Many other estimators of relatedness are available, including some based on the methods of moments [11, 28, 33-35]. We expect that adjustments similar to those proposed here for **G_1_** could be applied to these alternatives, but this would need to be investigated for each GRM.

We have shown that the numerator of the off-diagonal elements of **G_1_** has an expectation which does not depend on the depth, provided that, at a given depth, the probability of observing a single allele type (inferred as homozygous genotype) is the probability of that genotype inflated by a constant *K* (and conversely, the probability of observing a heterozygote is decreased by 2*K* from the true value). This includes the random sampling situation with *K* = 1/2 ^*k*^, where *k* is the depth. It also includes cases where observations of alleles are not independent, but are ‘clustered’, i.e. for a true heterozygote, the probability of a second read being the same as the first read is greater than 0.5. It is possible that this is actually the case, as random imbalances in allele numbers could be exacerbated during DNA amplification. In any case, we have shown that this will not affect the estimation of relatedness between two individuals.

On the other hand the diagonal elements of **G_1_** has an expectation which does depend on the depth, and we have shown how to correct this and demonstrated this correction in simulations using *K* = 1/2^*k*^ in both the simulation and the calculations. The application of this method to real data found self-relatedness estimates which, after some SNP filtering, did not appear to be related to the mean depth of the individual. More research with additional data sets at varying depths is needed to investigate whether there is clustering in allele reads, but this initial evaluation suggests that there is not a strong clustering effect.

We are hopeful that our methods will place less reliance on filtering data for low SNP call rates or low depth, as is common in the studies using GBS cited above. Our application of these methods to a real data set has demonstrated that SNP filters should be investigated, but to exclude data that deviate from the assumed model (e.g., of Mendelian inheritance). Atlantic salmon are known to have a genome which duplicated relatively recently, and still has regions of tetraploid inheritance [19]. This was evidenced by the ‘fin plot’ which showed many SNPs had near maximal heterozygosity for their given MAF. The filters we applied are just an illustration that appropriate filtering is likely to improve estimates. Optimal filters are likely to depend on the GBS methods (including differences in their mean depths) and on the species.

Although filtering for excess heterozygosity appears to have improved the estimation of relatedness, by giving results that better agree with the recoded pedigree, we still found estimates which appear to be low for highly related fish (full-sibs or parent-offspring). This could partly be due to incorrect assignment of samples to pedigree (either pedigree recording errors or sample recording errors). This is supported by the observation that some parents have very low estimated relatedness (near zero) with all or some of their putative progeny. Removing putative parent-offspring pairs where the mean estimated relatedness was less than 0.25 improved the mean estimated relatedness to 0.46 (analysis HW-.05sub). These possible mis-assignments are still under investigation, but it appears that the GBS results have been useful in identifying these issues. Another reason for depressed estimates of relatedness is genotyping error and regions with non-Mendelian inheritance. We have tried to reduce errors in the data by excluding low MAF SNPs at the bioinformatics step, however, there are likely to still be some errors. Further investigation is required to see if allowing for sequencing error can improve estimates. If it is not possible to eliminate the effects of error and non-Mendelian inheritance, one possibility would be to use estimates of relatedness that are regressed towards pedigree-derived values (after appropriate pedigree cleaning), along the lines of the third GRM method given by VanRaden [11].

Li *et al.* [36] argue that low depth (around 2-4) sequencing (allowing more individuals to be genotyped for the same cost) is a good strategy for GWAS. Gorjanc *et al.* [10] reached a similar conclusion for GS studies, finding that optimal prediction accuracy was obtained with low depth (around 1-2) sequencing of many individuals. These conclusions are in contrast to Pérez-Enciso [29], who found that a depth of 4 was too low, giving distorted GRMs. This situation should be ameliorated by using the methods proposed here. For between individual relationship estimation, our simulations suggest an optimal depth of 1-5 if the total sequencing effort is fixed (here the trade-off is between number of SNPs and SNP depth), i.e. low depth is desirable. Estimation of self-relatedness was optimised at a higher depth (5-10). This is because observation of both alleles of a SNP are needed for informing self-relatedness, whereas only a single observation is needed at a pair of individuals to inform between individual relatedness. The choice of depth for GBS studies will depend on the purpose of the study and what other resources are available. Studies which rely on allele frequency estimates of groups such as diversity studies are likely to benefit from lower sequencing depths and more individuals. However, in contrast it will be more difficult to estimate inbreeding with low coverage, but there may be other resources which can assist, e.g. pedigree information, or genomic methods that are based on runs of homozygosity [37] using SNPs with moderate coverage. The latter method requires a genomic map.

Our third simulation shows that it is mainly higher MAF SNPs that are contributing to the estimation of relatedness. Therefore it is likely to be beneficial to remove very low MAF SNPs (e.g. at the SNP calling step) as these may contain a high proportion of incorrect reads.

A possible drawback of our method is that it does not guarantee a GRM that is positive semi-definite, and so may not be invertable, which is a requirement for GBLUP. This will need investigation on a case by case basis. It may be possible to modify a non-invertible GRM to make it invertible [38] with only minor changes to the elements of the GRM. Another approach might be to take advantage of methods which require only a submatrix of the GRM to be inverted [39]. Individuals corresponding to that submatrix (e.g. the parents) could be sequenced at higher depth, while the other individuals (e.g. the selection candidates) could be sequenced at lower depth.

It may be possible to extend the approach to calculating GRM using GBS data outlined here to include calculations of linkage disequilibrium and linkage estimators. Such extensions or alternative approaches [40] would further enhance the utility of these low coverage sequencing approaches to genotyping. Currently sequencing is still relatively expensive so sequencing and analytical technologies that can sample restricted regions of the genome at low and variable depths but still provide accurate and unbiased data via simple methodology have a significant place for routine use to genetically improve agricultural species and for conservation genetics.

## Conclusions

We have addressed two major impediments to widespread use of genotyping by sequencing for routine use: the first is the ability to calculate unbiased estimates of the GRM using low coverage sequencing where genotypes are often missing and genotypes are probabilistic. The method makes maximal use of the available genotype data, is computationally efficient via use of matrix algebra and does not depend on problematic solutions such as imputation. Optimal sequencing depths for both relatedness and inbreeding were then calculated by simulation and the technique was also validated on a large real data set of pedigree recorded animals. The second impediment are SNPs with non-Mendelian inheritance based on observed GBS genotypes. A simple graphical method is given to illustrate this issue and to suggest an appropriate filter so they can be excluded from the GRM calculations. This works well for fixed differences between segmental duplications or allo-tetraploidy and also identifies SNPs that are segregating in one of the duplicated genome copies.

## Authors’ contributions

KGD, JCM and SMC conceived this study and provided guidance and oversight throughout the project. KGD developed the estimators used, undertook the simulation and statistical analysis of the salmon data. TK provided the salmon fin clips, data on the source of these samples and helped with interpretation of the results. SMC, RMA and TCvS processed the fin clips through to the creation of the sequencing results. RB undertook the bioinformatics analysis of the sequence results. All authors contributed to, read and approved the final manuscript.

## Acknowledgements

This project was supported by the Ministry of Business, Innovation and Employment via its funding of the “Genomics for Production & Security in a Biological Economy” programme (Contract ID C10X1306).

